# Acidic pH attenuates immune killing through inactivation of perforin

**DOI:** 10.1101/2024.07.04.602105

**Authors:** Adrian W Hodel, Jesse A Rudd-Schmidt, Tahereh Noori, Christopher J Lupton, Veronica CT Cheuk, Joseph A Trapani, Bart W Hoogenboom, Ilia Voskoboinik

## Abstract

Cytotoxic lymphocytes are pivotal effectors in our immune system. Their most potent way to delete virus infected or cancerous target cells is by perforin/granzyme dependent killing. Perforin is secreted into the immunological synapse and forms transmembrane pores in the target cell plasma membrane, allowing granzymes to enter the target cell cytosol and trigger apoptosis. The prowess of cytotoxic lymphocytes to efficiently eradicate target cells has been widely harnessed in immunotherapies against haematological cancers. Despite efforts to achieve a similar outcome against solid tumours, the immunosuppressive and acidic tumour microenvironment poses a persistent obstacle. Using different types of effector cells, including therapeutically relevant anti-CD19 CAR T cells, we demonstrate that the acidic pH typically found in solid tumours hinders the efficacy of immune therapies by impeding perforin pore formation within the immunological synapse. Nanometre-scale study of purified recombinant perforin undergoing oligomerization reveals that pore formation is inhibited specifically by preventing the formation of a transmembrane β-barrel. The absence of perforin pore formation directly prevents target cell death. This finding uncovers a novel layer of immune effector inhibition that must be considered in the development of effective immunotherapies for solid tumours.

**One Sentence Summary:** Our study reveals that an acidic environment hinders the efficacy of immune therapies by impeding the action of perforin, a critical protein in immune-mediated cell killing, directly within the immunological synapse.

## Introduction

Cytotoxic lymphocytes, encompassing cytotoxic T lymphocytes (CTLs) and natural killer (NK) cells, are immune effectors that rely on the pore-forming protein perforin and pro-apoptotic serine proteases, granzymes, to kill virus infected or transformed/pre-cancerous cells. Once activated, CTLs recognise their target and form an immunological synapse. Intracellular cytotoxic granules that store perforin and granzymes fuse with the plasma membrane, releasing their contents into the synaptic cleft. Once released, perforin forms large transmembrane pores in the plasma membrane of target cells [1], enabling diffusion of granzymes into the target cytosol where they trigger various apoptotic pathways [2].

In a common form of anti-cancer immunotherapy, T lymphocytes are genetically modified to express a chimeric antigen receptor (CAR), transforming them into CAR T cells that, following *ex vivo* expansion and re-infusion into the patient, direct their powerful cytotoxicity against cancerous cells. Currently, CAR T immunotherapies are successfully used against some haematological malignancies [3], but are largely ineffective against any solid tumours [4–6]. This is attributed to the immunosuppressive microenvironment of solid tumours [7, 8], prompting significant efforts to render it more permissive to immunotherapy. One aspect of immunosuppression that has received limited attention is the acidic environment of solid tumours. Tumour acidification occurs due to uptake of glucose and release of lactic acid into the tumour microenvironment as the preferred metabolic pathway [9]. Advances in magnetic resonance imaging using pH sensitive contrast agents provided visualizations of heterogeneous pH distributions *in vivo* in the extracellular milieu of solid tumours in mice [10–14] and humans [15], with pH frequently found between 6-6.5. Intratumoural pH below 6.5 has also been shown with a recently developed pH sensitive fluorescent cell surface marker [16].

The pore forming activity of perforin is exquisitely pH dependent, being most potent at neutral pH. Under these conditions, perforin monomers bind the lipid membrane in a calcium dependent fashion [17–19]. The monomers then assemble into small oligomers containing 2-7 protein subunits. As these oligomers are not yet inserted into the membrane, they are referred to as prepore assemblies or ‘prepores’. Initiated by their assembly and through a transition within the protein structure, prepores insert into the membrane and act as a nucleation site for further, rapid prepore binding and insertion, thereby growing the pore diameter [20]. Mature perforin pores are heterogeneous in diameter and curvature and can form arc- or ring-shaped structures [20–22]. In general, they contain 19-24 subunits and span lumen diameters of 10-20 nm [23, 24]. This process is greatly perturbed under acidic conditions: perforin lysis becomes reduced below pH 6.6 and is fully lost at pH 6 for both mouse [19, 25, 26] and human perforin [27]. At such low pH, calcium-dependent binding of perforin to lipid membranes remains intact, but the structural transitions necessary to form mature pores are blocked. Despite this, perforin is not denatured by the acidic pH and regains its lytic function upon restoration of pH levels to 7.4 [2, 27].

We hypothesized that inhibition of perforin pore formation at the acidic pH observed in solid tumours should limit immune killing. However, it has never been determined to what extent extracellular pH can affect perforin function within a physiological immunological synapse. Using different types of cytotoxic lymphocytes, including anti-CD19 CAR T cells, in a range of immunological assays, we assess how immune killing is influenced by pH. In addition, we use nanoscale imaging techniques to identify how perforin function is affected at acidic pH at a single-molecule level.

## Results

### Effector killing is attenuated at acidic pH despite sufficient release of perforin

To test the killing capacity of primary human immune effector cells, we used anti-CD19 CAR T cells that can readily recognize and kill CD19 positive target cells. As target cells, we transduced the U937 cell line, which exhibits a high sensitivity to CAR T cells, to express the extracellular domain of CD19 (hCD19t). To conduct assays at neutral pH 7.4 and acidic pH 6, we developed a pH-stable cell medium that does not require equilibration with CO_2_. To achieve this, we equilibrated bicarbonate free DMEM media with two buffers, 20 mM HEPES (pKa 7.3, 37 °C) and 20 mM MES (pKa 6, 37 °C) (**Supplementary Figure 1**).

To measure pH dependent immune killing, ^51^Cr-labeled U937-hCD19t cells were resuspended with anti-CD19 CAR T cells at different effector to target cell ratios and at various pH. The release of ^51^Cr from target cells after 4 h, where predominantly perforin/granzyme-mediated immune killing occurs, was gradually reduced with decreasing pH until it was abrogated at pH 6 (**Figure 1A**). The prolonged exposure to the acidic pH did not influence effector cell viability, and immune killing was fully restored after resuspending the cells to neutral pH (**Figure 1B**).

**Figure 1:**
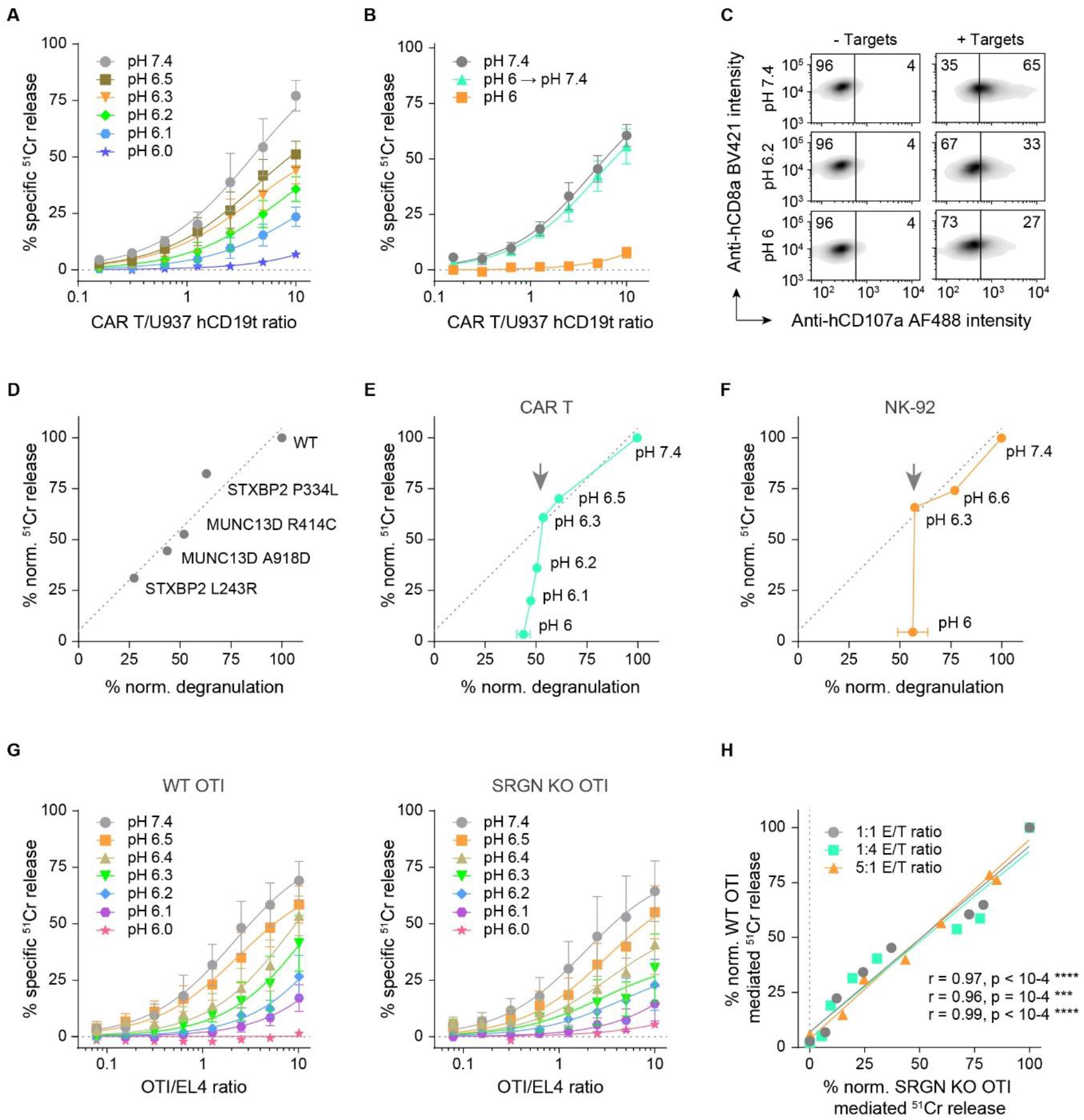
Immune killing and degranulation of different immune effectors. **(A)** Anti-hCD19 CAR T killing of U937 hCD19t targets at different pH, as a function of effector to target cell ratios, measured by the release of ^51^Cr radionuclide label. Data depicts mean ± standard error from three biological replicates. **(B)** Rescued immune killing of effectors that have previously been incubated at pH 6 for 4 h at 37 °C and were then resuspended at neutral pH and used in a killing assay immediately afterwards. Controls at pH 7.4 and pH 6 were using immune effectors without prior incubation at acidic pH. Data depicts mean ± standard error from three biological replicates. **(C)** Flow cytometry-based detection of degranulation of anti-hCD19 CAR T effectors in the absence (‘-Targets’) and presence (‘+ Targets’) of U937 hCD19t targets at neutral and acidic pH. Samples without targets were used to determine the cut-off for degranulation. Percentages of the CAR T cell population positive or negative for degranulation is indicated by the numbers in the upper corners. **(D)** Graph correlating degranulation and immune killing at neutral pH based on different hypomorphic mutations of granule trafficking proteins MUNC13D and STXBP2 obtained in a different study [30]. ^51^Cr immune killing was calculated from Michaelis-Menten fits of mean values from three biological replicates at a 1:1 effector to target ratio. Degranulation data shows mean of two biological replicates. Degranulation and ^51^Cr release were normalized to WT (100%). **(E)** Equivalent to D) but using data from anti-hCD19 CAR T killing and degranulation at different pH as shown in A) and C). The line fit is carried over from D). The arrow indicates the pH below which the data curve deviates from the line fit. Degranulation data shows mean ± standard deviation of biological triplicates (pH 7.4, pH 6) or a single experiment (pH 6.5 – pH 6.1). **(F)** As in D/E) using NK-92 effectors against K562 target cells. Degranulation data shows mean ± standard deviation of independent triplicate experiments (pH 7.4, pH 6) or a single experiment (pH 6.3, pH 6.6). **(G)** WT and SRGN KO OTI CTL killing of SIINFEKL pulsed EL4 cells at different pH. Data depicts mean ± standard error from three biological replicates. **(H)** Plot of WT vs SRGN KO OTI killing at indicated effector to target (E/T) ratios. Pearson’s correlation analysis with two tailed t-test for the different ratios is shown in the bottom right corner in the same order as the legend.

Since efficient CAR T cell killing requires sequential antigen binding, immune synapse formation and degranulation, perforin pore formation, and granzyme induced apoptosis, we reasoned that the disruption of any of these events would detriment CAR T cell cytotoxicity. Therefore, we first tested antigen binding and degranulation at acidic pH using externalisation of the granule associated protein CD107a (LAMP1) [28, 29]. We found that CAR T cell recognition of target cells, synapse formation and degranulation were progressively reduced, but not abolished by lowering media pH, with 27% of effector cells degranulating at pH 6, compared to 65% at pH 7.4 (58% reduction) (**Figure 1C**). Similar observations were made with the natural killer cell line NK-92, where no killing was observed at pH 6, despite an appreciable degranulation still being present (37% at pH 6 compared to 61% at pH 7.4; 39% reduction) (**Supplementary Figure 2**). To adequately interpret this apparent disparity between immune killing and degranulation, we next assessed whether a similar reduction in degranulation is sufficient to abrogate immune killing under more physiological conditions. To this end, we analysed recently published data on the effect of hypomorphic disease-causing mutations in two proteins responsible for cytotoxic granule exocytosis, MUNC13D and STXBP2, on CTL degranulation and cytotoxicity [30]. As expected, the degranulation and cytotoxicity of mutant CTLs were closely associated (**Figure 1D**). This contrasted with degranulation and cytotoxicity of CAR T and NK-92 cells at acidic pH, which exhibit a sharp drop in killing below pH 6.3 without a corresponding reduction in degranulation (**Figure 1E, F**).

We next evaluated whether reduced killing at acidic pH reflected reduced perforin release. Using a recently established experimental system [31, 32], we directly visualized ALFA-tag perforin (ALFA-PRF) secretion by murine CTLs [33] at pH 7.4 or pH 6.0, using an artificial immunological synapse formed between the CTL and glass coverslips coated with anti-CD3/CD28 antibodies. Fluorophore conjugated anti-ALFA-tag nanobodies were added to the culture media to visualize secreted protein, and the accumulating fluorescence within the artificial synapse quantified as shown in **Supplementary Figure 3**. Consistent with degranulation assays of CAR T effectors, we observed ∼50% reduced but still significant secretion of ALFA-PRF at pH 6.

Lastly, we assessed whether the released perforin could still be sequestered by the cytotoxic granule storage protein serglycin. The interaction between serglycin and perforin is physiologically important [34] and pH-dependent [35], in that perforin and serglycin are hypothesised to dissociate at the neutral pH of the immunological synapse. Having confirmed that the acidic pH had no effect on SIINFEKL antigen presentation by EL4 target cells (**Supplementary Figure 4**) and that target cell killing by transgenic OT1 CTLs recovered at pH 7.4 after their prolonged exposure to the media with pH 6 (**Supplementary Figure 5**), we then determined the activity of wild-type and serglycin-deficient OTI CTLs at different pH (**Figure 1G**). We found that the two cell types responded to the changing pH identically (**Figure 1H**) suggesting that serglycin plays no role in attenuating immune killing at acidic pH.

### pH dependent immune killing correlates with pore forming activity of perforin

As our live-cell data supported the idea that acidic extracellular pH affects perforin function after its release into the immunological synapse, we assessed the molecular basis of pH-dependent pore formation by perforin. To this end, we used purified recombinant murine wild-type perforin (WT-PRF) in conjunction with turbidity-based sheep red blood cell (SRBC) lysis assays (**Supplementary Figure 6A**,**B**) and atomic force microscopy (AFM) on solid-supported phospholipid bilayers. MMT buffer (see Methods) had no detectable effect on the lytic activity of WT-PRF compared to the commonly used buffer but controlled the experimental conditions at pH 4-9 (**Supplementary Figure 6C**). SRBCs were stable overnight at pH ≥ 5.5 (**Supplementary Figure 6D**), and a supported lipid bilayer as used in AFM experiments remained stable at pH ≥ 4 for the duration of the experiments (∼1 h).

We set up SRBC lysis assays at a range of WT-PRF concentrations and the pH range between 5.5-7.5. The change of turbidity that reflected SRBC lysis was monitored over 6 h at 37 °C. The results highlight that acidification slows cell lysis and concomitantly reduces the maximum achievable level of lysis (**Figure 2A**). Extreme concentrations of WT-PRF were still able to lyse SRBCs at pH 6 (at reduced capacity), but not at 5.5. In addition, molecular scale images from AFM experiments (for practical reasons at a single WT-PRF concentration and not time-resolved) produced a marked reduction of WT-PRF arc- and ring-shaped pores at pH 6-6.5 and their complete absence at and below pH 5.5 (**Figure 2B**). Quantification of WT-PRF pore formation and subsequent fit with a four-parameter logistic curve yielded a half maximum inhibition at pH 6.4 ± 0.2 (mean ± 95% confidence interval) (**Figure 2C, black**). These findings were remarkably consistent with SRBC lysis data (**Figure 2C, orange**), and immune killing (**Figure 2C, blue**), directly linking the attenuation of immune killing to reduced WT-PRF pore formation at acidic pH.

**Figure 2:**
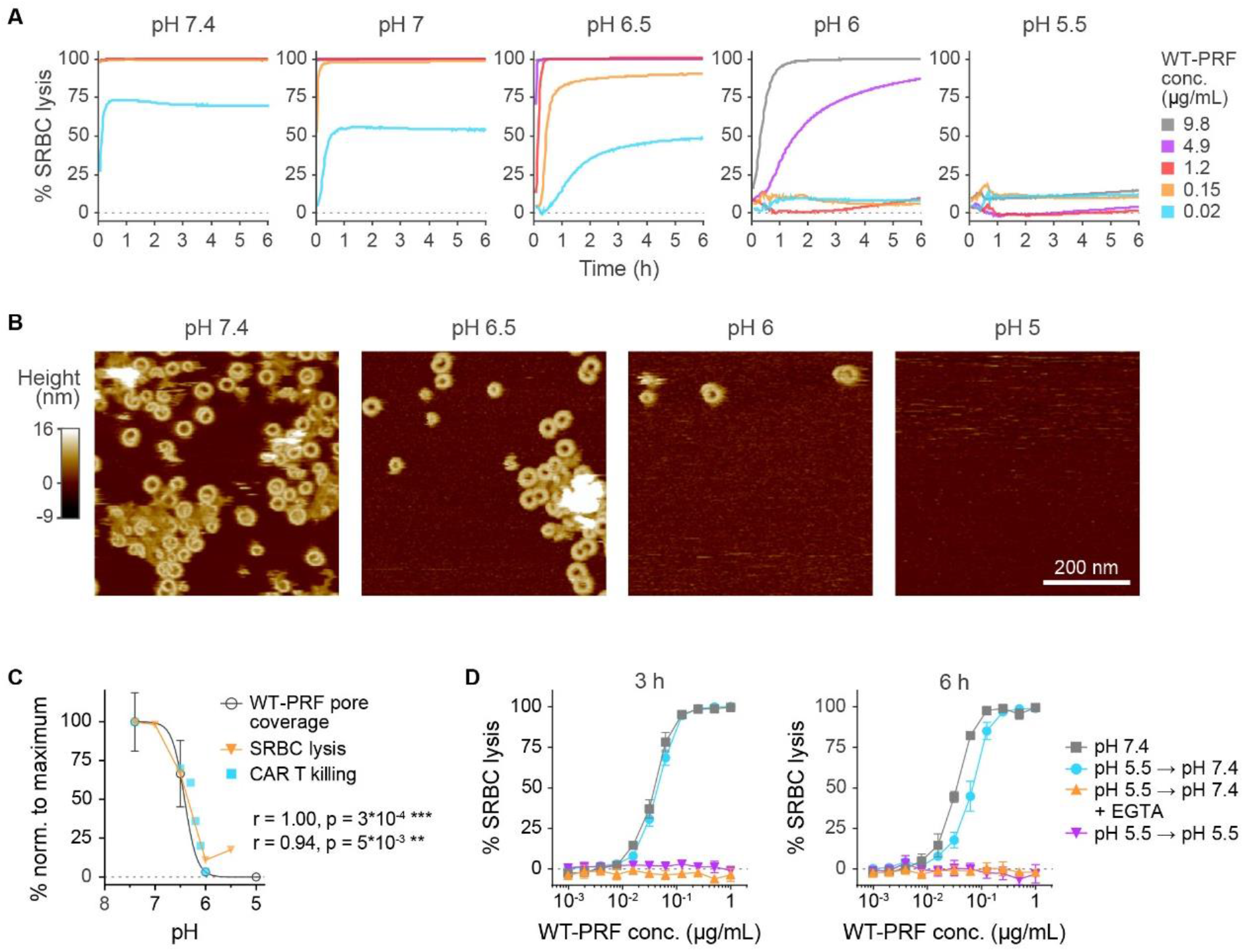
WT-PRF mediated lysis and pore formation is reduced at acidic pH and can be restored. **(A)** 6 h timelapse of WT-PRF mediated lysis of SRBCs at pH 7.4-5.5 and five different protein concentrations at 37 °C. 1 µg/mL ≈ 16 nM WT-PRF. **(B)** AFM images of supported lipid bilayers incubated with WT-PRF at the indicated pH. Perforin pores are visible as golden ring- and arc-shaped features, while the lipid background is coloured maroon. Pore formation is gradually reduced with pH, and below pH 6, no pore features were observed. The images shown are representative of five different areas assessed across the sample surface. **(C)** Overlay of SRBC lysis shown in A) at 0.15 µg/mL and 0.5 h, WT-PRF pore coverage quantified from AFM data as in B), and CAR T cell immune killing at a 1:1 effector to target ratio from **Figure 1A**. All values were normalized to lysis, pore formation, and immune killing, respectively, observed at pH 7.4 (100%). WT-PRF pore coverage was fitted with a four-parameter logistic curve. The fitted values were used to perform Pearson’s correlation analysis with two-tailed t-test against SRBC lysis or CAR T killing data, respectively; corresponding correlation coefficients are shown in the bottom right side of the graph. **(D)** Recovery of SRBC lysis after incubation at pH 5.5 and indicated incubation times (3 h or 6 h) when neutralizing pH. The calcium chelator EGTA was added while restoring pH in control samples. Data shows mean ± standard error from three independent experiments.

WT-PRF function in the presence of SRBCs and Ca^2+^ ions was largely restored, even after hours-long incubation at pH 5.5, by neutralizing pH (**Figure 2D**). Similarly, AFM images show recovery of arc- and ring-shaped pore assemblies after neutralizing pH on samples that initially exhibited few or no pores (**Supplementary Figure 7A**). The size distribution of these assemblies represents an endpoint characteristic for a pore forming pathway. We therefore traced perforin assemblies in the collected AFM data (see Methods) of samples that have been recovered from acidic pH and compared the resulting size distributions with pores formed at neutral pH (**Supplementary Figure 7B**). The shape of these size distributions was conserved across all pH levels, thus indicating that WT-PRF at acidic pH may initially interact with the lipid membrane normally, and after pH is restored, recommences the natural pore forming pathway. Following the already well-established mechanism of perforin pore formation [20], we went on to explore how acidic pH inhibits the perforin pore.

### Perforin remains in prepore-like assemblies at acidic pH

Perforin is released into the extracellular space as soluble monomer. Using mass photometry [36] to measure the molecular weight of WT-PRF in solution (**Supplementary Figure 8A, B**), we found that it exists as a monomer at neutral pH but, at pH 5.5, ∼70% of the protein forms dimers and ∼20% remains in a monomeric form, with the rest of the protein found in higher order assemblies (**Supplementary Figure 8C**). Restoring the pH from 5.5 to 7.4 causes dissociation back into monomeric protein (**Supplementary Figure 8D**).

We next assessed perforin binding to lipid membranes at acidic pH. Using AFM, we can detect superficially bound WT-PRF at pH 5 after trapping it into plaques by adding a crosslinker (**Supplementary Figure 9**). The height of these plaques corresponds to the height of an upstanding perforin monomer, ∼11 nm [20], in line with a membrane-proximal location of the lipid binding C2 domain of perforin [18, 23]. Since C2 domain binding is calcium-dependent [17, 19], we further tested if that is also the case at acidic pH using SRBC binding assays in the presence or absence of 1 mM Ca^2+^ and at neutral pH or pH 5.5. To prevent cell disintegration during the assays at neutral pH, we used a non-lytic murine perforin mutant with an engineered disulphide-lock, TMH1-PRF [20]. The flow cytometry data (**Figure 3A**) shows membrane binding of TMH1-PRF at pH 7.4 in the presence of Ca^2+^ and some binding in the absence of Ca^2+^. Presumably, the latter one does not involve the essential C2 domain, since the unlocked TMH1-PRF (by reduction of its engineered disulphide bond) has no haemolytic activity under these conditions (**Supplementary Figure 10**). At pH 5.5, TMH1-PRF binding only occurs in the presence of Ca^2+^. Taken together this data suggests that perforin binds the lipid substrate at acidic pH through its C2 domain, as it would under neutral conditions.

**Figure 3:**
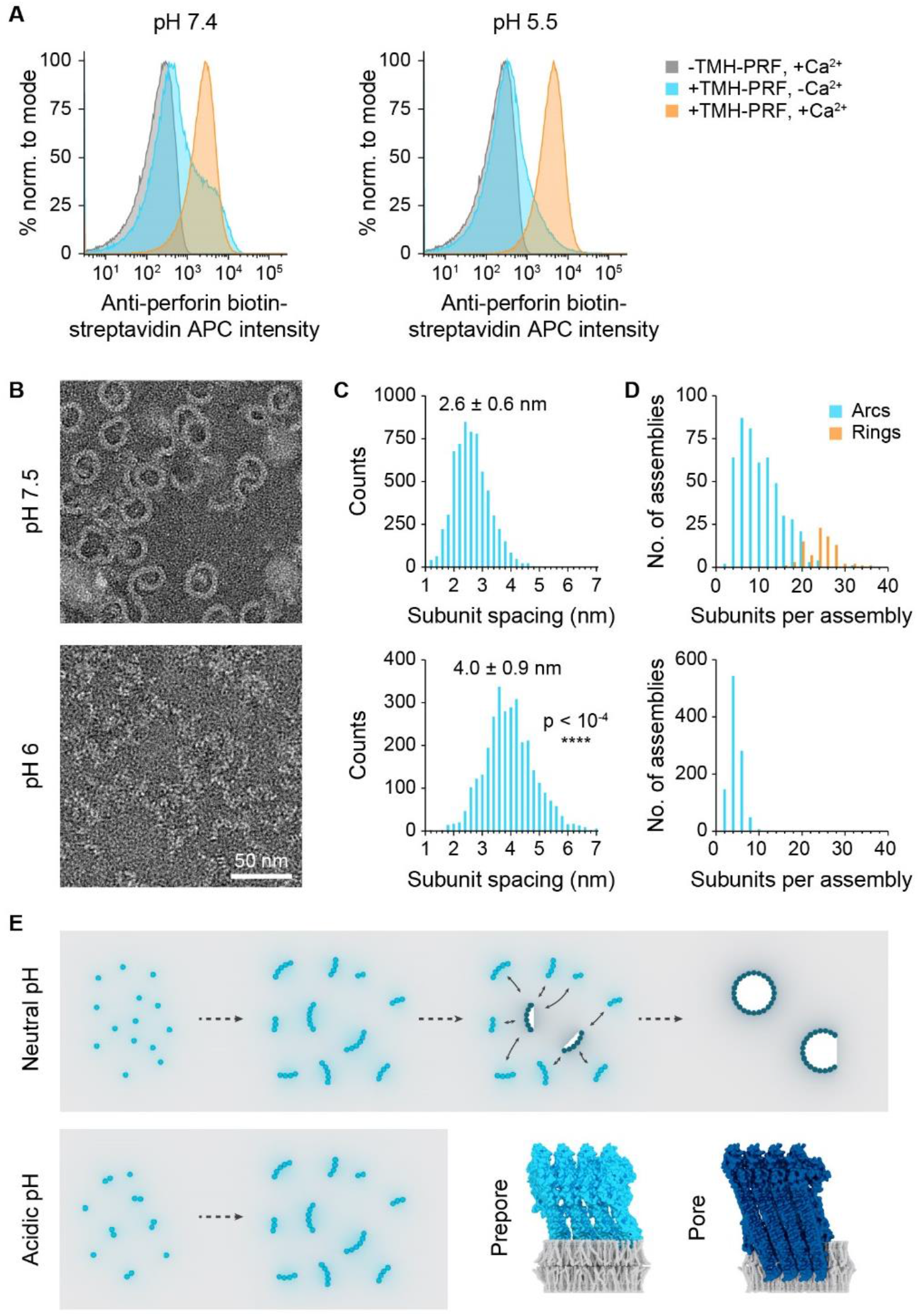
EM images of perforin bound to lipid monolayers show prepore-like assemblies at acidic pH. **(A)** Binding of 1 µg/mL non-lytic perforin mutant, TMH1-PRF, in the presence and absence of 1 mM Ca^2+^ and at pH 7.4 and 5.5 against an untreated control at pH 7.4, evaluated by flow cytometry. **(B)** EM images of lipid monolayers containing WT-PRF assemblies formed at the indicated pH levels. **(C)** Histograms of spacings between adjacent WT-PRF subunits extracted from EM images. Mean ± standard deviation of subunit spacing is indicated atop each histogram. Distributions were significantly different in Kolmogorov-Smirnov test of unbinned and χ^2^ test of binned data, both p < 10^−4^ (two-tailed) as indicated in the lower histogram. Number of analysed subunits were n ≈ 6000 (pH 7.4) and n ≈ 3000 (pH 6). **(D)** Histograms of subunits per WT-PRF assemblies shown in A). Ring- and arc-shaped assemblies are plotted separately in orange and blue, respectively. Number of analysed assemblies were n = 495 (pH 7.4) and n = 1028 (pH 6). A total sample area of 2.7 µm^2^ was analysed for each pH in C) and D). **(E)** Schematic illustration of the molecular assembly pathway of perforin pore formation at neutral pH, compared to acidic pH. The two schematics on the bottom right illustrate the concept of membrane surface bound non-lytic prepores and membrane inserted, lytic pores. Perforin models were used from the RCSB protein data base (rcsb.org), accessions 3NSJ and 7PAG.

Once perforin binds to the lipid membrane, it assembles into short, linear prepores that typically contain only two to five protein subunits [20, 24]. They are thus much smaller than mature pores that typically contain more than 16 subunits and can close into a ring-shape. In addition, average spacing between subunits changes from 3.9 nm in prepores to 2.6 nm in pores [20]. With this information, we sought to identify which part of the perforin assembly pathway was affected by acidic pH. We used a negative-stain electron microscopy dataset [2] of perforin bound to lipid monolayers at pH 6 and at neutral pH in MMT buffer (**Figure 3B**) and quantified it as follows: the subunit spacing at pH 6 was measured at 4.0 ± 0.9 nm compared to 2.6 ± 0.6 nm (mean ± standard deviation) at pH 7.4 (**Figure 3C**). Furthermore, the assemblies formed at pH 6 contain fewer than ten protein subunits throughout, in contrast to the larger arc- and ring-shaped assemblies formed at pH 7.4 (**Figure 3D**). By comparing these results with the behaviour of perforin prepores and pores, we conclude that at acidic pH, perforin is unable to insert its transmembrane helices to line a β-barrel pore and remains in a prepore-like state, as illustrated in **Figure 3E**.

We finally considered the known structures of perforin [23, 24] to develop a hypothesis for the structural basis of this behaviour at acidic pH. Perforin pore formation depends on the unfurling of two α-helical bundles per molecule to result in two transmembrane β-hairpins. These hairpins connect with each other and with hairpins of adjacent subunits via hydrogen bonds, forming a stable β-barrel structure lining the pore [24]. Intriguingly, the intra- and inter-subunit interface between hairpins of perforin is lined by seven (murine perforin) or eight (human perforin) histidine residues with a sidechain pKa ≈ 6 (**Figure 4**). Acidification would protonate these sidechains, reducing the number of possible hydrogen bonds between them. The functional implication of such weaker intermolecular binding would be a less efficient unfurling of the hairpins, hence reduced pore formation and protection against cytotoxic lymphocyte killing as demonstrated in our data.

**Figure 4:**
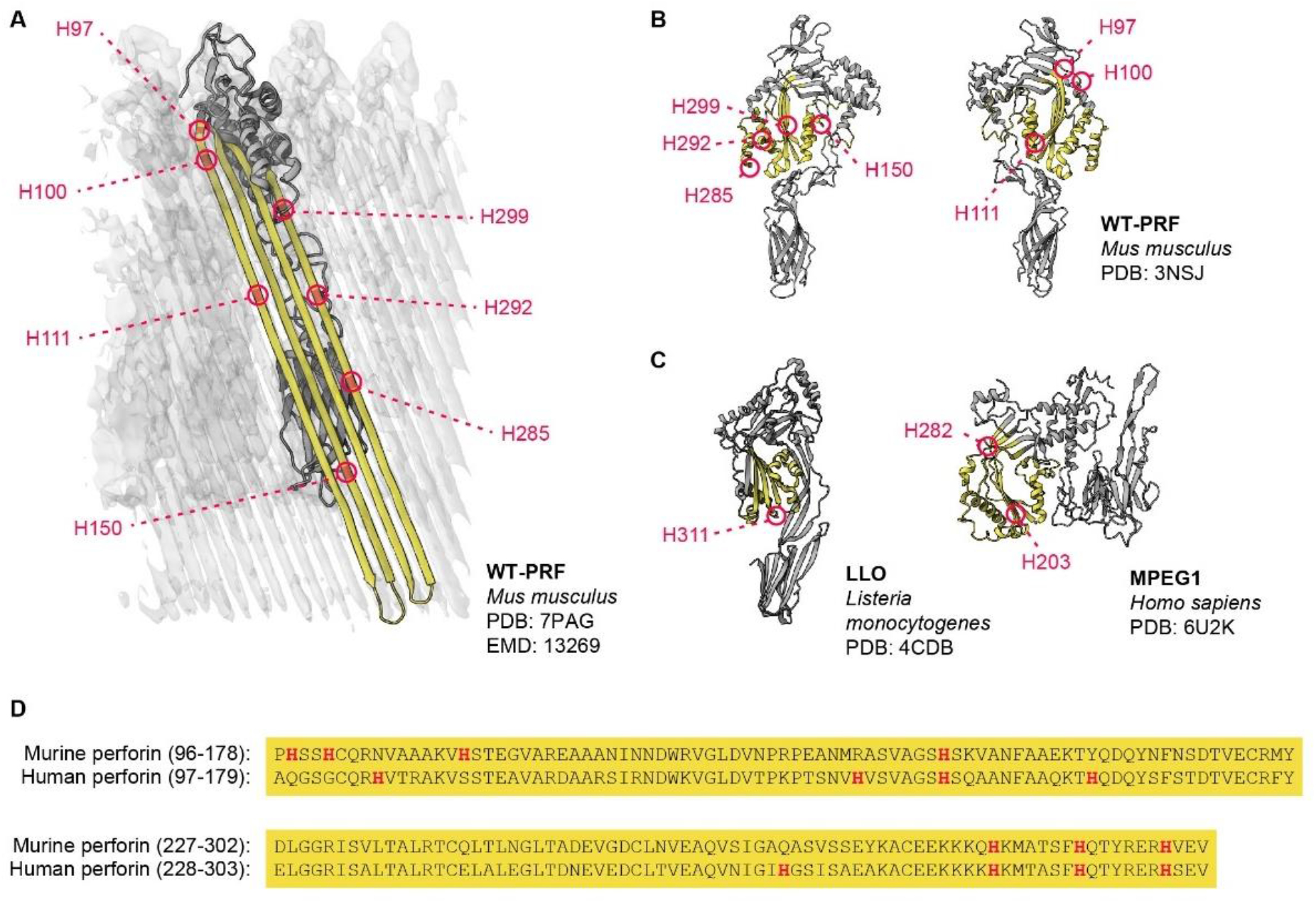
Histidine residues at transmembrane β-hairpin interfaces of perforin. **(A)** Cartoon of a (murine) WT-PRF subunit in the pore state, overlayed with a section of an electron density map of the perforin pore (grey). The transmembrane β-barrel (yellow) motif of WT-PRF is seamed with numerous histidine residues (red). **(B)** The same histidine residues as in A) highlighted in the soluble monomer structure of WT-PRF, shown from two sides. **(C)** By comparison, the acid activated pore forming proteins listeriolysin O (LLO) and macrophage expressed gene 1 (MPEG1) contain fewer histidine residues (red) in the corresponding regions (yellow, green). Protein models were generated from the RCSB protein data base using the indicated model accessions. **(D)** Comparison between wild-type murine and human perforin amino acid sequence for the β-barrel motif. The sequences were retrieved from UniProt (uniport.org) under accessions P10820 and P14222, respectively, and aligned. Histidine residues are highlighted in red and are similarly abundant in both murine and human PRF, but not necessarily conserved.

## Discussion

We tested the effect of extracellular acidification on the killing ability of various types of cytotoxic lymphocytes - primary human CAR T cells, an NK cell line, and primary mouse CTLs. Despite having fundamentally different receptor-antigen interactions, they all showed similar sensitivity to acidic pH, resulting in the rapid decline of immune killing at pH lower than 6.5, and the loss of activity at pH 6.

Between pH 7.4 and pH 6.5, the attenuation of cytotoxicity correlates with the degree of lymphocyte degranulation. This was not surprising, given our own experimental data and other reports on degranulation inhibitors in CTLs [37] and granule trafficking defects in immunodeficiencies [38]. The finding is also in line with the pH sensitivity of ORAI1/STIM1 channels that control calcium influx and degranulation [39] and operate at approximately half capacity at pH 6.5 compared to neutral pH [40]. However, below pH 6.5, immune killing was acutely reduced and then lost at pH 6. There, we also observed the reduction in degranulation, but this could not sufficiently explain the complete loss of function of cytotoxic lymphocytes. Instead, we found a near perfect correlation between immune killing and the ability to of perforin to form transmembrane pores. Inactivation of perforin was thus the critical underlying cause of reduced immune killing below pH 6.5.

The importance of our work is reflected by the persistently low efficacy of CAR T cell therapies against solid tumours. Within these tumours, the microenvironment acts as an unyielding barrier. Factors like checkpoint signalling and diminished cell metabolism [7, 8, 41–43] hinder the long-term performance of immune effectors in eliminating their targets. However, potent immune killing ultimately hinges on perforin mediated target cell death. Addressing perforin inactivation at the acidic intratumoural pH is thus necessary to improve clinical outcomes of immunotherapy.

The challenge of perforin inactivation at acidic pH can possibly be solved by raising the pH of the tumour microenvironment [44, 45], which would also reduce other immunosuppressive effects. Alternatively, immune cells could be engineered to withstand acidity and maintain their cytotoxicity. To this end, our molecular analysis of how acidic pH impacts perforin provides a guide to the design of pH insensitive perforin.

## Materials and Methods

### Antibodies

A full list of antibodies used in this study is provided in **Supplementary Table 1**. Unless otherwise denoted, 2∗10^5^ nucleated cells or ∼2∗10^7^ sheep red blood cells (SRBCs) were stained in 50 μL of cell culture media or MMT buffer (pH 7.4) containing antibody diluted 1:100 for 30 min on ice. Primary/secondary stains were incubated sequentially, and the stained cells resuspended in 100-200 µL of cell culture media or buffer solution for flow cytometry.

### Cell culture

Primary human PBMCs and CAR T derivatives thereof were cultured in RPMI 1640 (Gibco, Life Technologies, Carlsbad, CA, USA) with 10% FCS, 2 mM Glutamax, 10 mM HEPES, 1 mM sodium pyruvate, 100 μM non-essential amino acids, 50 IU/mL penicillin, 50 μg/mL streptomycin, and 100 IU/mL recombinant human IL2 (National Cancer Institute, Washington, Maryland, USA). To induce proliferation of CTLs after harvest, 30 ng/mL of anti CD3 antibody clone OKT3 was added to the media once.

Primary mouse splenocytes from C57BL/6.OTI transgenic mice were cultured in RPMI 1640 with 10% FCS, 2 mM Glutamax, 10 mM HEPES, 1 mM sodium pyruvate, 100 μM non-essential amino acids, 50 IU/mL penicillin, 50 μg/mL streptomycin, 50 μM 2-mercaptoethanol (CAS 60-24-2, Sigma-Aldrich), and 100 IU/mL recombinant human IL2. To induce proliferation of splenocytes after harvest, 10 ng/mL of SIINFEKL peptide were added to the media once. Murine experiments are approved by the Peter MacCallum Cancer Centre Animal Ethics Committee (AEEC) E655.

EL4 cells were cultured in DMEM (with high glucose, L-glutamine, and phenol red) with 10% FCS and 2 mM Glutamax. U937 cells were cultured in RPMI 1640 with 10% FCS and 2 mM Glutamax. Cells were pulsed with 1 µg/mL SIINFEKL peptide (Genscript, Piscataway, NJ, USA) for 1 h at 37 °C prior to use as targets for mouse OTI CTLs.

Cells in RPMI 1640 were cultured at 37 °C, 5% CO_2_, and cells in DMEM at 37 °C, 10% CO_2_.

### Anti-hCD19 CAR T cells

CAR T effectors were generated from activated PBMCs isolated from healthy human donor blood. The cells were transduced with anti-hCD19 CAR using either retroviral or lentiviral transduction. For retroviral transductions, activated PBMCs were transduced with anti-hCD19 (clone FMC63) scFv_myc_hCD8hCD28_hCD3**ζ** CAR subcloned into a pSAMEN vector backbone using the supernatant of stably retrovirus producing PG13 cells and Retronectin (Takara Bio, Kusatsu, Shiga, Japan) coated culture plates as per manufacturer’s instructions. For lentiviral transductions, activated PBMCs were exposed to the viral supernatant of HEK293T cells transfected with anti-hCD19 (clone FMC63) scFv_flag_hCD8hCD28_hCD3**ζ** CAR subcloned into a pUltra vector backbone, and third generation lentivirus packaging vector in the presence of 0.25% v/v Lentiboost (Sirion Biotech, Gräfelfing, Germany). To obtain a pure population of anti-hCD19 CAR T cells, transduced PBMCs were sorted for CD8^+^/CD16^-^/myc^+^ or CD8^+^/CD16^-^/flag^+^ using fluorescently activated cell sorting (FACS) 6-7 days after harvest. CAR T cells were used up to 20 days after harvest.

### U937 hCD19t target cell line

For hCD19 CAR T effectors to produce the highest readouts in degranulation assays, we empirically selected and designed the U937 hCD19t target cell line. To generate a sequence for human CD19t with truncated intracellular signalling domain (hCD19t), the open reading frame from the full length hCD19 nucleotide sequence (GenBank NM_001770) was truncated at base pair 969, analogous to published work [46]. The sequence was subcloned into an MSCV GFP vector backbone (Addgene 91975). HEK293T cells were transfected with hCD19t MSCV GFP and ampho retroviral packaging vector. The retroviral supernatant was used to transduce U937 cells in Retronectin coated plates as per manufacturer instructions. The transfected cells were sorted twice with respect to GFP reporter levels and anti-hCD19 antibody binding using FACS.

### pH stable cell media

pH adjusted media for live cell assays was prepared from powdered, bicarbonate-free DMEM powder (Gibco) dissolved in Milli-Q water at the manufacturer specified concentration and supplemented with 0.1% fatty acid free bovine serum albumin (Roche Diagnostics, Mannheim, Germany), 1X Glutamax, 20 mM HEPES, and 20 mM MES hydrate (CAS 1266615-59-1, Sigma-Aldrich). The pH was subsequently adjusted to the desired levels at 37 °C by addition of 1 M NaOH, monitored by a calibrated pH meter. Media toxicity was assessed as shown in **Supplementary Figure 1** and was found to be comparable to standard T cell medium. pH stable cell media osmolarity was assessed by the Peter MacCallum Cancer Centre Media Kitchen facility and ranged between 300-320 mOsmol/kg, i.e., between T cell and standard DMEM culture media (**Supplementary Table 2**).

### ^51^Cr release assays

As a measure of cell lysis, target cells were labelled with 50-100 µCi ^51^Cr radionuclide as sodium chromate in normal saline (PerkinElmer, Waltham, MA, USA) and incubated at 37 °C, 5% CO_2_ for 1 h. Effector and target cells were washed and resuspended in pH buffered media adjusted to the appropriate pH. A titration series of effector cells was combined with target cells in a 96 well plate containing 10^4^ targets and 10^5^ effectors in 200 µL media per well at the highest effector to target ratio. Wells containing either no effectors or 5% triton-X solvent solution were set up in parallel for each pH level as controls for spontaneous and total ^51^Cr release, respectively. The plates were incubated at 37 °C, 10% CO_2_ for 4 h. The remaining cells were pelleted afterwards, and the supernatant extracted for γ-radiation decay counting on a Wizard2 Gamma Counter (PerkinElmer). The percentage of specific ^51^Cr release in a well *n* was calculated using the radiation counts per minute (CPM): Specific ^51^Cr release (%) = 100 **/** (CPM_total lysis_ – CPM_spontaneous lysis_) ∗ (CPM_n_ – CPM_spontaneous lysis_). Michaelis-Menten kinetics were fitted with the maximum set to V ≤ 100% and the Michaelis constant set to k_M_ > 0 as boundary conditions.

### Degranulation assays of anti-CD19 CAR T cells

Degranulation was measured by detecting CD107a on the surface of effector cells. To this end, 5∗10^4^ effector cells were mixed with 2∗10^5^ target cells in 200 µL of media and incubated for 3 h. Since antibody affinity changes with pH, all samples were washed three times with media at neutral pH before labelling in 50 µL of RPMI (pH 7.4) containing 1:100 anti-hCD8 BV421 and 1:25 anti-hCD107a AF488 antibodies for 30 min on ice. Samples were washed once after incubation and then resuspended in 100 µL of media for flow cytometry.

### Expression and purification of WT-PRF and TMH1-PRF

Recombinant wild-type mouse perforin (WT-PRF) and disulphide locked mutant (A144C-W373C) perforin (TMH1-PRF) were expressed and purified using baculovirus infected Sf21 cells as described previously [47]. The protein was eluted in 270 mM imidazole, 300 mM NaCl, 20 mM Tris, pH 7-8 at concentrations between 0.1-0.5 mg/mL.

### MMT buffer

Buffers were prepared in 0.9% saline solution (154 mM NaCl). To produce MMT buffered saline, 10 mM of DL-malic acid (CAS 6915-15-7, Sigma-Aldrich, St. Louis, MO, USA), 20 mM of MES hydrate, and 20 mM of Tris (CAS 77-86-1, Merck Millipore, Darmstadt, Germany) were added and adjusted between pH 4-8.5 using sodium hydroxide or hydrochloric acid. 1-5 mM of CaCl_2_·2H_2_O (CAS 10035-04-8, Sigma-Aldrich) or 4 mM EGTA (CAS 67-42-5, Sigma-Aldrich) were added to buffers as outlined in the text.

### SRBC cell lysis timelapse assays

An SRBC stock was kept in Celpresol (Immulab, Parkville, VIC, Australia) at 4 °C. Prior to each assay, SRBCs were washed and resuspended at ∼2∗10^8^ cells/mL in MMT buffer adjusted to the desired pH levels and containing 2 mM Ca^2+^. WT-PRF dilution series were set up on ice in 100 μL of buffer (without Ca^2+^) and combined with 100 μL of SRBCs per well of a flat-bottom 96-well plate. Wells containing no WT-PRF were used as spontaneous lysis controls, and SRBCs lysed in Milli-Q water were used in total lysis controls. As a measure of lysis, turbidity was monitored in 2-minute intervals at 600 nm absorption using a Cytation 3 plate reader (BioTek, Agilent Technologies, Santa Barbara, CA, USA) preheated to 37 °C and after equilibrating for 20 min. Lysis percentage in a well *n* was calculated using the absorbance values *A*: Lysis (%) = 100 **/** (A_total lysis_ – A_spontaneous lysis_) ∗ (A_n_ – A_spontaneous lysis_).

### Atomic force microscopy

AFM sample preparation and imaging largely followed previously published methods [22]. In brief, 4 µL of a 1 mg/mL suspension of unilamellar vesicles made from DOPC (Avanti Polar Lipids, Alabaster, AL, USA) were deposited on freshly cleaved mica (∼10 mm diameter) covered in 100 µL MMT buffer, 25 mM MgCl_2_, 5 mM CaCl_2_, pH 7.4 and incubated for 30 min at room temperature until a supported lipid bilayer was formed. To adjust pH, the samples were washed 9 times with 80 µL MMT buffer, 25 mM MgCl_2_, 5 mM CaCl_2_, pH 4-6.5. Subsequently, WT-PRF was injected at a final concentration of ∼150 nM and incubated for 5 min at 37 °C, and then imaged by AFM. To restore pH to neutral, the samples were washed a further 9 times with 80 µL MMT buffer 25 mM MgCl_2_, 5 mM CaCl_2_, pH 7.4, incubated for 15 min at 37 °C, and re-imaged. AFM images were recorded on a MultiMode 8 system (Bruker, Santa Barbara, CA, USA) using PeakForce Tapping at 2 kHz with MSNL-E and -F probes (Bruker) at 50-100 pN of force. Images were processed in the NanoScope Analysis software (version 1.8, Bruker) for background flattening and colour (height) scale adjustments with the supported lipid bilayer surface set to 0 nm.

### SRBC lysis recovery from acidic pH

WT-PRF dilution series were set up in 50 µL of MMT at pH 7.4 and pH 5.5 without Ca^2+^ on ice and combined with 25 µL of dilute SRBCs (∼6∗10^8^ cells/mL) in MMT containing 3 mM Ca^2+^ per well of a 96-well flat-bottom plate. For SRBCs at pH 7.4, 125 µL of MMT pH 7.4, 1 mM Ca^2+^ were added immediately, and the plate was incubated for 3 h or 6 h at 37 °C. To recover pH 5.5 to neutral pH, 125 μL of MMT pH 8.5, 1 mM Ca^2+^, with or without 4 mM EGTA was added per well, gently mixed, and incubated for a further 30 min at 37 °C. To maintain pH 5.5, 125 µL of MMT pH 5.5, 1 mM Ca^2+^ were added instead. For total lysis controls, 125 µL of water were added instead of buffer. Absorption of each well at 600 nm was measured on a Cytation 3 plate reader, and lysis calculated as outlined before.

### Antibody based detection of purified TMH1-PRF

TMH1-PRF was incubated with SRBCs at a concentration of 1 μg/mL for 15 min on ice under the desired conditions. Cells were washed once and resuspended in RPMI media at neutral pH for antibody staining, as outlined under ‘Antibodies’. The cross-species reactivity and use of anti-human perforin clone δG9 to detect disulphide locked murine perforin mutant TMH1-PRF was evaluated in **Supplementary Figure 11**.

### Electron microscopy data analysis

Negative-stain electron microscopy data from an earlier study [2] was generously provided by Helen Saibil and Natalya Lukoyanova of Birkbeck College, London, United Kingdom. The data was analysed on a user interface written in-house in Matlab (MathWorks, Natick, MA, USA) as used previously [22]. The generally well-resolved subunits of perforin assemblies in electron microscopy data allowed for manual marking of subunits to extract the number of subunits per assembly and inter-subunit distance, determined by linear interpolation.

## Supplementary Materials

Supplementary Methods

Supplementary Figure 1: pH stable cell media tests for cell viability and pH stability.

Supplementary Figure 2: NK-92 cells degranulate but do not kill K562 target cells at acidic pH.

Supplementary Figure 3: Release of ALFA-PRF into artificial synapses at different pH.

Supplementary Figure 4: SIINFEKL is presented by H-2Kb (MHC I) at pH 6.

Supplementary Figure 5: OTI immune killing of SIINFEKL pulsed EL4 target cells recovers after neutralizing pH.

Supplementary Figure 6: Evaluation of SRBC lysis assays using turbidity in MMT buffer.

Supplementary Figure 7: AFM detection of WT-PRF pores at different pH before and after neutralization.

Supplementary Figure 8: Mass photometry of WT-PRF in solution.

Supplementary Figure 9: WT-PRF binds a supported lipid bilayer at pH 5 as detected by AFM.

Supplementary Figure 10: TMH1-PRF has no lytic activity in the absence of Ca^2+^.

Supplementary Figure 11: Anti-human perforin antibody reacts with disulphide-locked TMH1-PRF.

Supplementary Figure 12: Analysis strategies for flow cytometry data.

Supplementary Figure 13: FACS strategy for CAR T cells.

Supplementary Table 1: Table of antibodies used in this study.

Supplementary Table 2: Osmolarity measurements of pH stable media compared to standard cell media.

Supplementary Table 3: STR results and analysis for cell line identification.

## Acknowledgements

We thank Helen Saibil and Natalya Lukoyanova (Birkbeck, University of London) for kindly providing electron microscopy data, for supporting this study, and critical reading of the manuscript; Annette Ciccone and Sandra Verschoor (Peter MacCallum Cancer Centre) for preparing purified perforin; Andrew Ellisdon (Monash University) for support of mass photometry experiments, Neil Young (University of Melbourne) for providing sheep red blood cells; Jane Oliaro (Peter MacCallum Cancer Centre) for providing the NK-92 cell line. Experiments were supported by the Peter MacCallum Cancer Centre Flow Cytometry and Genomics Cores, Media Kitchen, and Centre for Advanced Histology and Microscopy.

## Funding

AWH – Swiss National Science Foundation p2skp3_187634 and P500PB_211089.

BWH and IV – UK Medical Research Council MR/V009702/1.

IV – Cancer Council Victoria and NHMRC (2011020).

## Author contributions

AWH – designed the study, designed, conducted, and analysed experiments, co-wrote manuscript.

JRS – designed, conducted, and analysed experiments, read and commented on manuscript.

TN – designed, conducted, and analysed experiments, read and commented on manuscript.

CJL – designed, conducted, and analysed experiments, read and commented on manuscript.

VCC – conducted and analysed experiments, read and commented on manuscript.

JAT – co-lead research, read and commented on manuscript.

BWH – co-lead research, read and commented on manuscript.

IV – designed the study, designed experiments, lead research, co-wrote manuscript.

## Data and materials availability

All data needed to evaluate the conclusions in the paper are present in the paper or the Supplementary Materials.

