## Supplementary Methods, Figures, and Tables for "Acidic pH attenuates immune killing through inactivation of perforin"

***Flow cytometry***

Flow cytometry data for degranulation of effector cells and perforin detection on SRBCs was analysed as shown in **Supplementary Figure 12**. Flow cytometry based fluorescently activated cell sorting of CAR T cells was carried out by the Peter MacCallum Cancer Centre Flow Cytometry Core Facility using a gating strategy as outlined in **Supplementary Figure 13**.

***Cell line verification***

Human derived target cell lines U937 and K562 have been verified by small tandem repeat (STR) analysis shown in **Supplementary Table 3**. STR profiles were recorded by the Australian Genome Research Facility (AGRF, Melbourne, Victoria, Australia). NK-92 effector cells were verified by flow cytometry-based phenotype analysis showing NKp44$^{+}$, Cd56$^{+}$, CD54$^{+}$, CD45RA$^{+}$, and CD16$^{-}$ expression (**Supplementary Figure 2D**, CD16 not shown) in line with literature [48, 49]. Mouse derived EL4 target cells were verified by flow cytometry-based analysis of the mCD90.2 phenotype found 100% positive compared to a human control cell line (data not shown). Cells were tested regularly for mycoplasma contamination by the Peter MacCallum Cancer Centre Genomics Core.

***Mass photometry of perforin in solution***

Stock solutions of WT-PRF or bovine serum albumin (BSA, Roche) in elution buffer, MMT pH 7.4, or MMT pH 5.5 at a concentration of 5 µg/mL were prepared. MMT buffers contained 1 mM $CaCl_2$. Mass photometry measurements were performed on a TwoMP system (Refeyn, Oxford, United Kingdom). The system was first focused with 16 µL of the respective buffer and subsequently injected with 4 µL of the stock solution to reach a final concentration of 1 µg/mL and the molecular weight data recorded. To restore WT-PRF in MMT pH 5.5 to neutral pH, MMT pH 8.5 was added to triple the sample volume. The system was then focused with 8 µL of buffer, and 12 µL of the restored sample was added to reach a final concentration of 1 µL/mL. All preparation and measurements were performed at room temperature.

***Viability of effector cells at pH 6***

To assess the viability of OTI/CAR-T effectors during our experimental conditions, $10^6$ effectors were stained with 200 µCi of $^{51}$Cr, washed, and resuspended at a concentration of $10^6$ cells/mL in 200 µL of bicarbonate free DMEM, 20 mM Hepes, 20 mM MES, BSA or FCS as indicated, pH 7.4 and pH 6. For reference, mouse/human T cell media at pH 7.4 or at pH 6 (acidified with HCl) with MES added as indicated was used. CAR-T effectors have additionally been exposed to U937 hCD19t effectors at a concentration of $10^6$ cells/mL (1:1 E/T ratio). After 4 h incubation at 37 °C and levels of $CO_2$ as indicated, cells were pelleted and 100 µL of supernatant extracted for radiation detection.

***PH stability of cell media***

2 mL of different media was aliquoted into 10 mL centrifuge tubes. pH levels of media were measured before and after incubation for 4 h at 37 °C at levels of $CO_2$ and with or without effector/target cells as indicated. Where cells were present, effector cells were used at a density of $10^6$ cells/mL and target cells at $10^5$ cells/mL respectively, producing a 10:1 E/T ratio at the same cell densities used in immune killing assays.

***WT-PRF fixation in AFM samples***

Samples containing WT-PRF assemblies on a DOPC bilayer formed at pH 5 were incubated with 0.2% glutaraldehyde 8% solution (TAAB Laboratories Equipment, Aldermaston, Berks, United Kingdom) for 6 h at room temperature and subsequently washed three times with 80 µL MMT buffer, 25 mM $MgCl_2$, 5 mM $CaCl_2$, pH 5 before re-imaging.

***K562 cell culture***

K562 cells were cultured in RPMI 1640 with 10% FCS, and 2 mM Glutamax and incubated at 37 °C, 5% $CO_2$.

***SRBC lysis measurements using haem release***

As an alternative to turbidity-based measurements, SRBC lysis can be calculated based on colorimetric measurement of haem released by the ruptured cells. For these measurements, killing assays were set up the same way (see Methods) in 96-well plates. After the assay, the plate is centrifuged to pellet SRBCs, and 100 µL of supernatant is transferred to a flat bottom 96-well plate. The absorbance *A* of each well at 410 nm was measured on a Cytation 3 plate reader and the lytic activity in a well *n* calculated as: SRBC lysis (%) = $100/(A_{total\ lysis} - A_{spontaneous\ lysis}) * (A_n - A_{spontaneous\ lysis})$.

***Fluorescence microscopy of artificial immune synapses using ALFA-PRF OTI CTLs***

Our experiments followed published protocols [32]. In brief, OTI CTLs were transduced with ALFA-PRF Tag-BFP MSCV and Lifeact-eGFP MSCV to visualize F-actin and sorted for Tag-BFP/eGFP double positive cells using FACS. Ibidi µ-Slide 18 well 1.5H glass bottom chamber wells (Ibidi, Martinsried, Germany) were coated with 10 µg/ml anti-mCD3ε and 5 µg/ml anti-mCD28 antibodies in PBS overnight at 4 °C. The wells were washed with phosphate buffered saline before use and equilibrated at 37 °C. Immediately prior to the experiment PBS was replaced by pH stable media at pH 6 or pH 7.4 containing ~$10^5$ transduced OTI CTLs and 50 nM FluoTag-X2 anti-ALFA AZDye 568 nanobodies (Nanotag Biotechnologies, Göttingen, Germany). The release of ALFA-PRF over time was monitored with a Zeiss Elyra PS1 microscope equipped with an alpha Plan-Apochromat 100x oil lens (both Zeiss, Oberkochen, Germany) in total internal reflection fluorescence (TIRF) mode. In the recorded images, cell boundaries were traced manually using the F-actin signal to extract the area where ALFA-PRF is detected therein. Manual tracing was performed with samples blinded.

***NK-92 cell culture and assays***

NK-92 cells were cultured in RPMI 1640 with 10% FCS, 2 mM Glutamax, 10 mM HEPES, 1 mM sodium pyruvate, 100 µM non-essential amino acids, 50 IU/mL penicillin, 50 µg/mL streptomycin, and 200 IU/mL recombinant human IL2. pH-sensitive experiments were carried out in RPMI 1640 with 10% FCS, 2 mM Glutamax, 10 mM HEPES, 1 mM sodium pyruvate, 100 µM non-essential amino acids, 50 IU/mL penicillin, 50 µg/mL streptomycin, 20 mM MES as buffer, besides sodium bicarbonate present in RPMI 1640. PH levels were adjusted by dropwise addition of 1 M malic acid at 37 °C. Note that this medium does not maintain pH values as stably as the pH stable medium used for CAR T cells, and pH values may rise by up to 0.4 increments during the assay. Killing and degranulation assays using NK-92 effector cells against K562 target cells were otherwise carried out as with CAR T cells.

#### Supplementary Figures

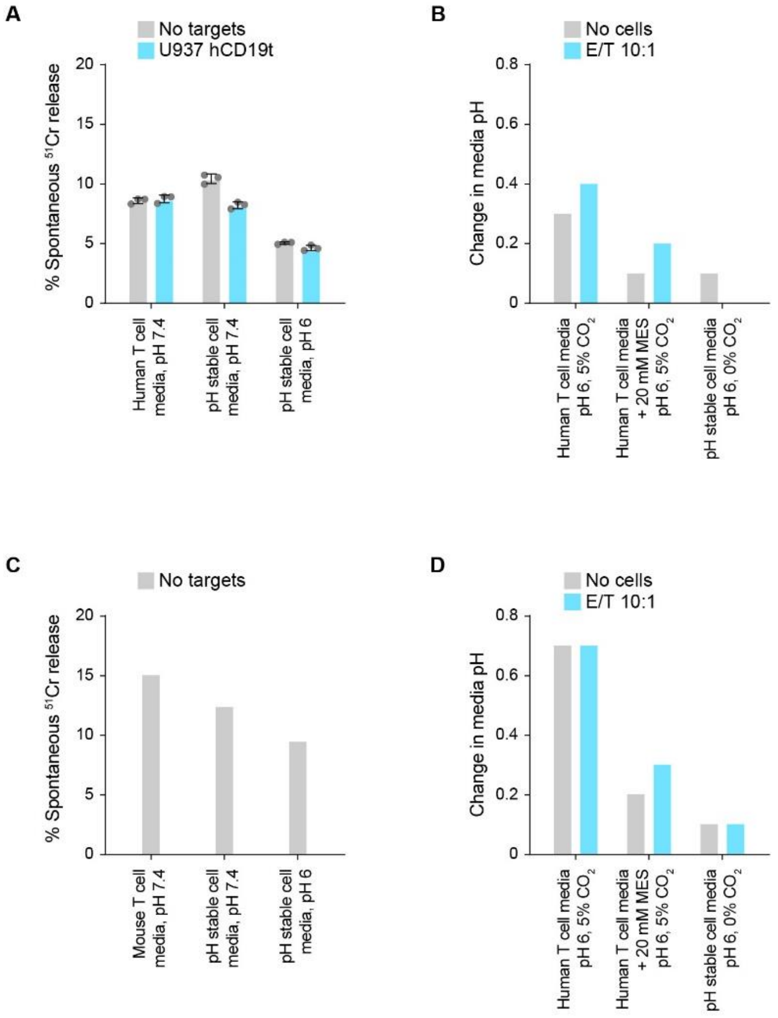


**Supplementary Figure 1: pH stable cell media tests for cell viability and pH stability.**
**(A)** To confirm that effector cell viability in pH stable media is comparable to human T cell media, spontaneous $^{51}$Cr release of anti-hCD19 CAR T cells was assessed after 4 h at 37 °C, 0% $CO_2$ in pH stable cell media at pH 7.4 and pH 6, or 5% $CO_2$ in human T cell media at pH 7.4. Cell density was at $5*10^5$ CAR T cells/mL, with or without $5*10^4$ U937 hCD19t target cells/mL.
**(B)** To assess the stability of pH 6 in different media, we measured the pH before and after 4 h at 37 °C and plotted the increase in pH. Media pH stability was assessed either without cells or in the presence of $5*10^5$ CAR T effectors and $5*10^4$ U937 hCD19t targets (10:1 E/T ratio).
**(C)** Analogous measurement of cell viability as in A) using mouse derived OTI effectors.
**(D)** Analogous measurement of pH stability as in B) using either no cells or OTI effectors and SIINFEKL pulsed EL4 target cells.
Measurements in A) represents mean ± standard deviation of three technical replicate samples, B-D) represent values from single experiments. pH values were rounded to the closest 0.1 increment. Overall, pH stable media supports similar effector cell viability as a standard culture media and have a stable pH (0.1 pH increment) under the conditions used in our assays.

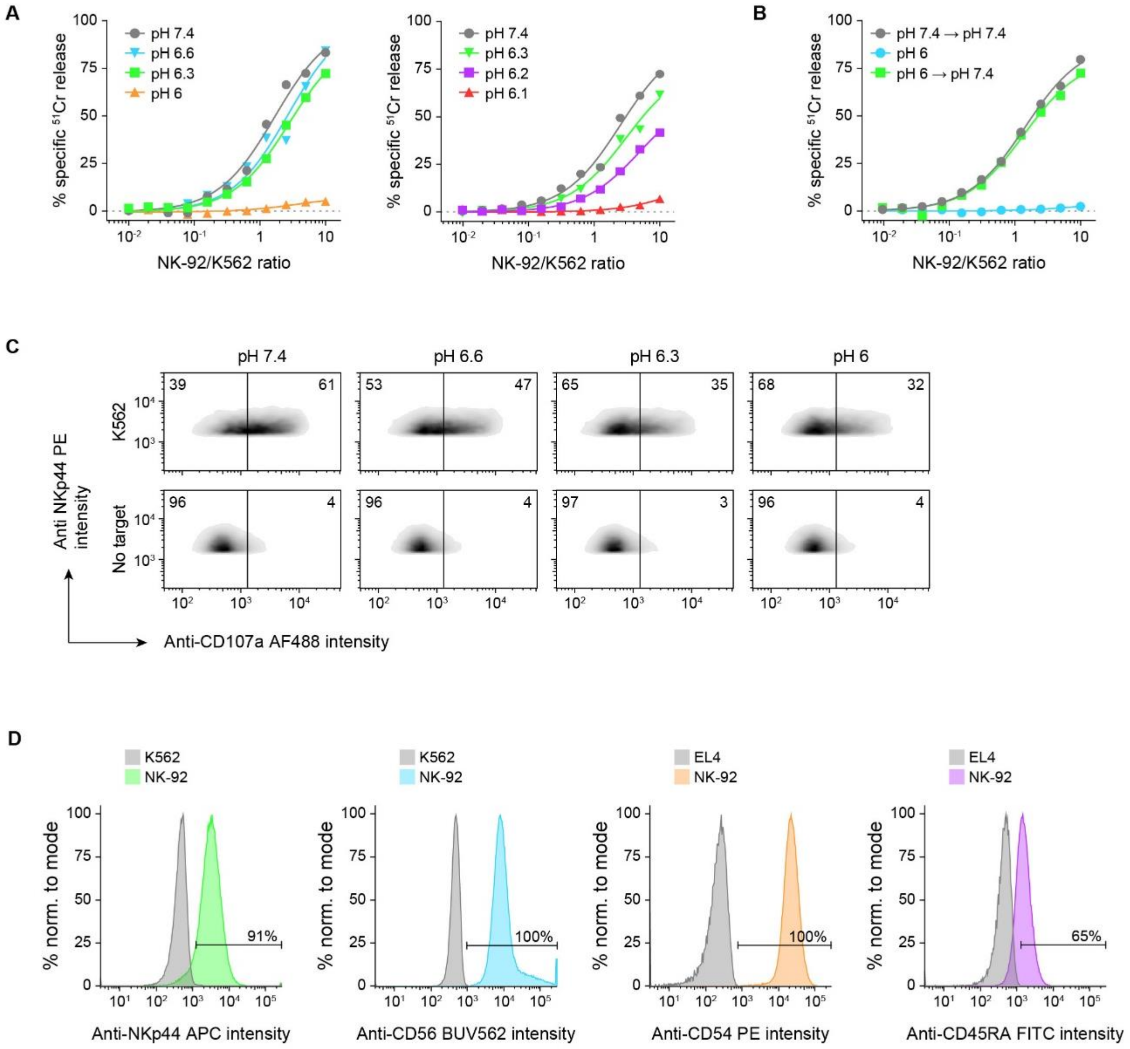


**Supplementary Figure 2: NK-92 cells degranulate but do not kill K562 target cells at acidic pH.**
**(A)** Killing assays performed at different pH outline a marked reduction of killing between pH 6.3-6.
**(B)** Immune killing of cells incubated for 4hr at pH 6 and 37 °C was fully restored after overnight incubation at neutral pH, 37 °C.
**(C)** Degranulation of NK-92 cells after mixing with K562 target cells at different pH shows a reduction, but not abrogation of degranulation. Of note, degranulation at pH 6.3 and 6 was almost identical, while the killing capacity was abolished (see A).
**(D)** Flow-cytometry based phenotype analysis of NK-92 for cell line verification.
All data was recorded in single experiments.

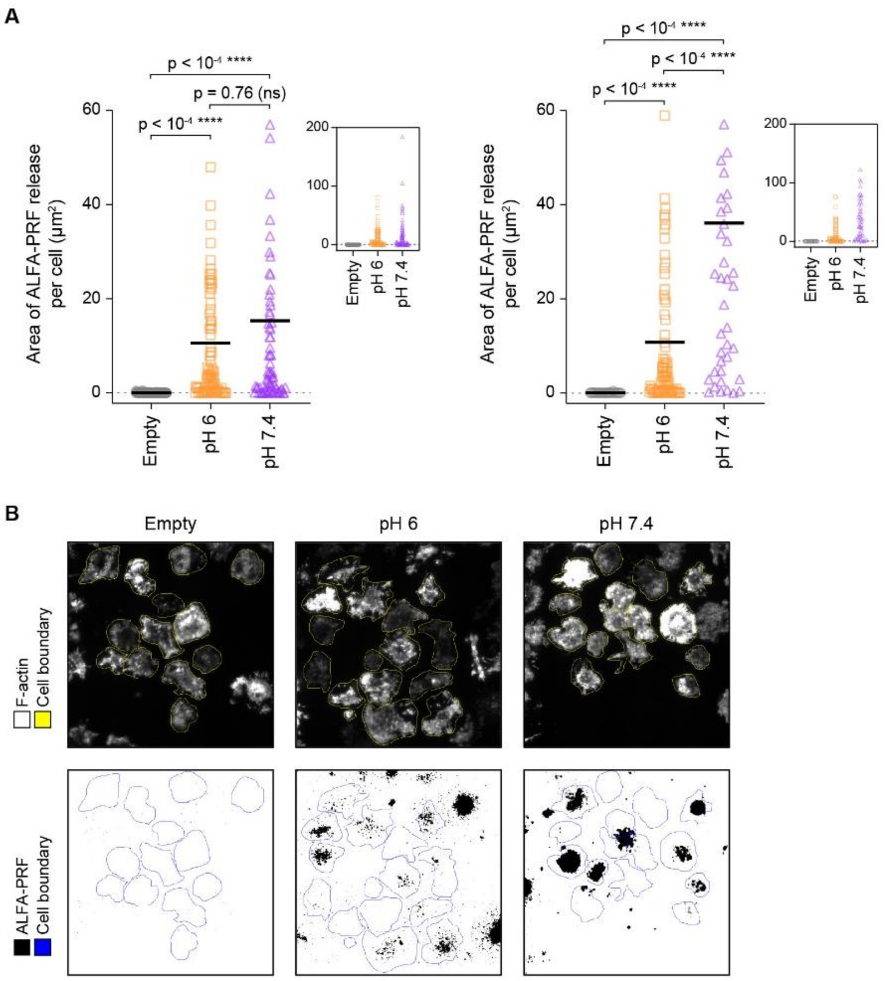


**Supplementary Figure 3: Release of ALFA-PRF into artificial synapses at different pH.** **(A)** Data from two biological replicates showing the area of ALFA-PRF release per cell into the artificial synapse. Cells that were not expressing ALFA-PRF (‘Empty’, shown in grey) were used as negative control and compared to ALFA-PRF release at pH 6 (orange) and pH 7.4 (purple). Data is shown magnified (large plots) or at full range (insets). Horizontal lines depict arithmetical means. The significance levels are determined using the p-value from Kruskal-Wallis and uncorrected Dunn’s post-hoc tests. **(B)** Representative fluorescence images of actin (top) and ALFA-PRF (bottom) displayed by effector cells engaged in artificial synapses. ALFA-PRF images are shown in binary after background subtraction. The coloured contours are manually traced cell boundaries used to calculate the area of ALFA-PRF release per cell. All experiments were blinded.

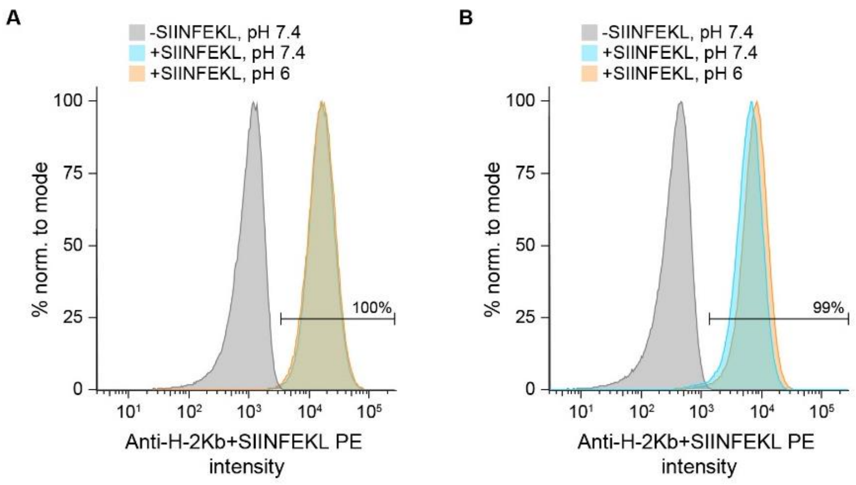


**Supplementary Figure 4: SIINFEKL is presented by H-2Kb (MHC I) at pH 6.**
**(A)** EL4 cells were pulsed with SIINFEKL at pH 7.4 and subsequently incubated at pH 6 for 4 h at 37 °C. The cells were then stained with anti-H-2Kb+SIINFEKL PE at neutral pH and their fluorescence measured by flow cytometry.
**(B)** EL4 cells were pulsed with SIINFEKL at pH 6 and then stained with anti-H-2Kb+SIINFEKL PE at neutral pH. The percentage gate is shown for populations incubated at pH 6, compared to untreated EL4 cells.

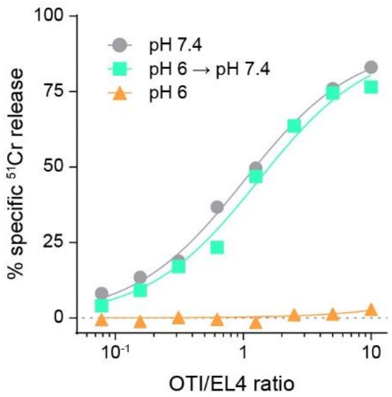


**Supplementary Figure 5: OTI immune killing of SIINFEKL pulsed EL4 target cells recovers after neutralizing pH.**

OTI immune cells were incubated for 4 h in pH stable media at pH 6, 37 °C, 0% $CO_2$, after which the pH was neutralized and an immune killing assay was set up immediately, including controls for OTI effectors that have not been in acidic conditions prior, at pH 6 and pH 7.4. After neutralizing pH, OTI effectors killed their EL4 targets at the same rate as without prior acidification. Data represents mean from n = 2 biological replicates.

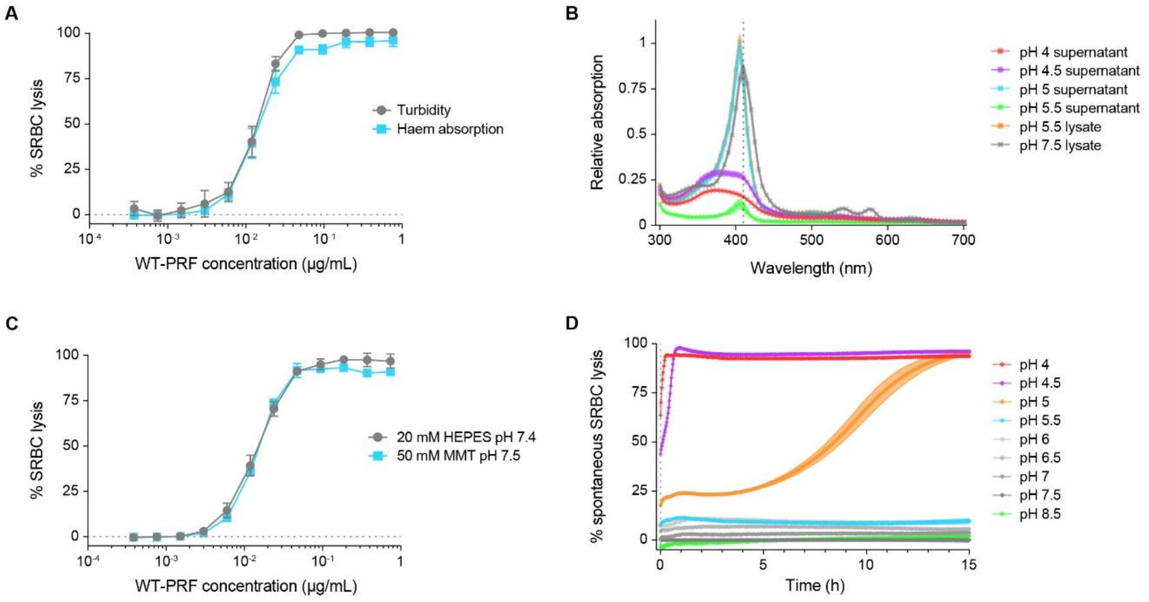


**Supplementary Figure 6: Evaluation of SRBC lysis assays using turbidity in MMT buffer.**
**(A)** SRBC lysis curves produced by WT-PRF show the same concentration dependent behaviour when using haem release or change in turbidity to detect lysis.
**(B)** Absorption spectra of haem at different pH, recorded between 300-700 nm in 5 nm increments of supernatant or pellets lysed in water as denoted, after 15 h incubation at 37 °C. Acidification produced a shift in the haem absorption peak to lower wavelengths (410 to 405 nm) accompanied by a ~20% increase of absorption at pH 5.5 and pH 5, and the loss of a defined peak at pH 4.5 and pH 4. This renders haem absorption less suitable to measure perforin dependent lysis at acidic pH, compared to turbidity-based measurements.
**(C)** Using turbidity change as detection method, WT-PRF induces identical concentration dependent SRBC lysis in HEPES and in MMT based buffers at neutral pH.
**(D)** In a timelapse measurement of spontaneous SRBC lysis at different pH using turbidity change as detection method, SRBCs remained stable for 15 h at pH 5.5-8.5 and 37 °C, and spontaneously lysed at pH 4-5.
The data shown in A) and C) was collected from three independent experiments and depicts mean ± standard error. The data shown in B) and D) was collected from three technical replicates and shows mean ± standard deviation as coloured background band.

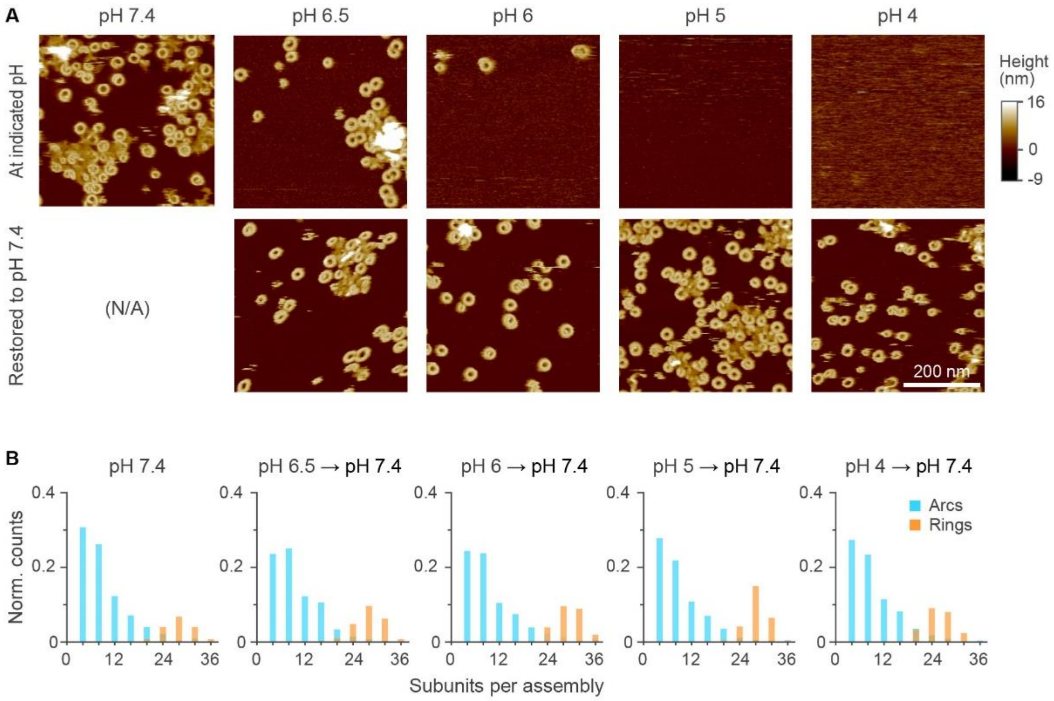


**Supplementary Figure 7: AFM detection of WT-PRF pores at different pH before and after neutralization.**

**(A)** The top row of AFM images exemplifies pH dependent WT-PRF pore formation. The samples were incubated with WT-PRF for 5 min at 37 °C and at indicated pH levels and subsequently imaged at room temperature for up to 1 h. The bottom row shows the same samples after restoring the pH by washing with pH 7.4 buffer and incubating for 15 min at 37 °C. The number of pores is visibly reduced at pH 6 before restoration and is absent in samples incubated at pH 5 and 4. After restoration of pH, WT-PRF pores emerge again in all samples, irrespective of prior pH. N/A, not assessed.

**(B)** Assembly size distributions of WT-PRF pores after restoration appear similar between pH levels, indicating that WT-PRF can recommence its pore formation using the same oligomerization pathway as in neutral pH. The data shown in B) was collected from 5 images taken across the sample surface, covering a total area of 1.5 $\mu m^2$ each.

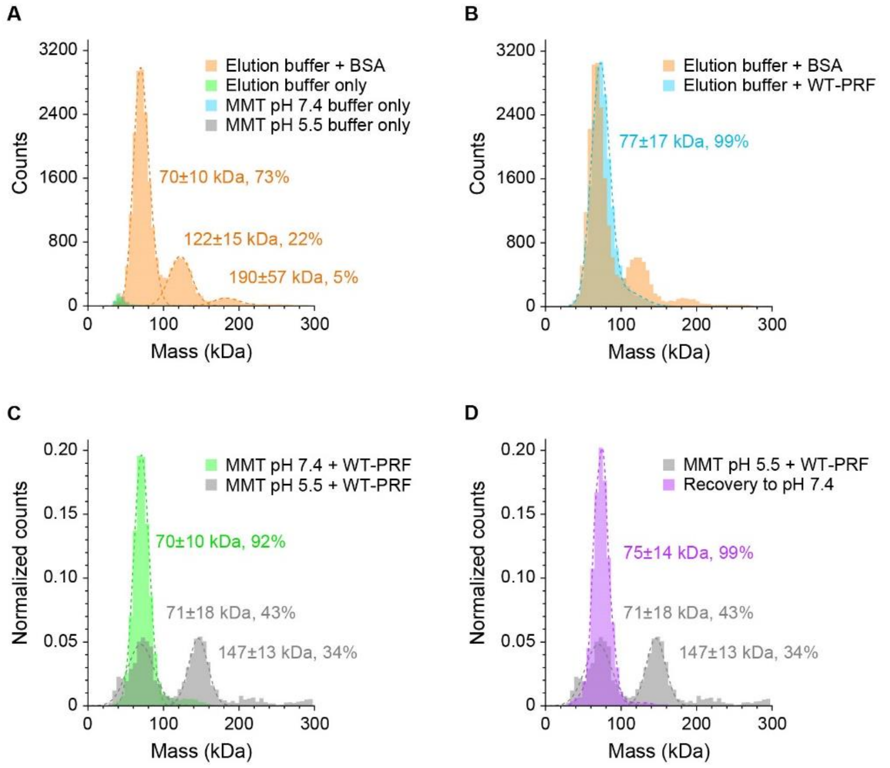


**Supplementary Figure 8: Mass photometry of WT-PRF in solution.**
**(A)** The prominent monomer and dimer peaks of bovine serum albumin (BSA, orange) in elution buffer were used to calibrate the mass photometry system. The buffers alone (green, blue, grey) produced a minor signal at around 40 kDa, which we interpret as noise at the lower boundary of the detection range.
**(B)** Wild-type murine perforin (WT-PRF, blue) in elution buffer produced a single peak overlapping with the BSA (orange) monomer signal and estimated at 77 kDa molecular weight, approximately corresponding to monomeric WT -PRF.
**(C)** A similar singular peak for monomeric WT-PRF is observed in MMT buffer at pH 7.4 (green), whereas at pH 5.5, two peaks with molecular masses approximately corresponding to monomeric and dimeric WT-PRF are dominant (grey).
**(D)** Dimeric WT-PRF found in MMT at pH 5.5 (grey, same as in C) disassembled into monomers upon restoring the buffer to pH 7.4 (purple).

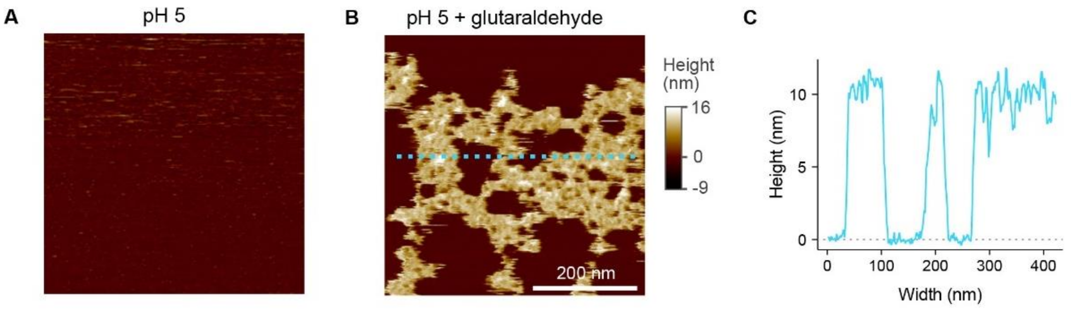


**Supplementary Figure 9: WT-PRF binds a supported lipid bilayer at pH 5 as detected by AFM.**
**(A)** At pH 5, no WT-PRF pores are detected by AFM in the supported lipid bilayer. However, potentially bound protein that is not membrane inserted is not readily resolved by AFM due to its high lateral mobility.
**(B)** To assess whether WT-PRF binds the lipid membrane at pH 5 but is too mobile, we added glutaraldehyde to cross-link bound protein into larger plaques and slowing it sufficiently to be resolved. At pH 5, we indeed observe the formation of plaques after addition of glutaraldehyde, indicating that WT-PRF binds at pH 5 without forming pores.
**(C)** These plaques reach 11 nm in height as determined by a cross-section taken across the dashed line in B), corresponding to the height of an upstanding perforin monomer.

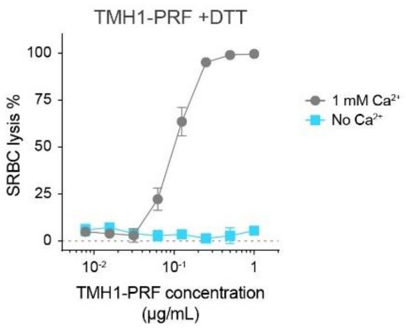


**Supplementary Figure 10: TMH1-PRF has no lytic activity in the absence of $Ca^{2+}$.** The disulphide locked mutant TMH1-PRF was titrated and incubated with SRBCs in MMT pH 7.4 in the presence of 4 mM of the reducing agent DTT, which reduces the TMH1 disulphide bond and restores lytic function, and with or without 1 mM $Ca^{2+}$. Lysis was detected as a measure of change in turbidity and plotted against TMH1-PRF concentration, showing lytic activity exclusively in the presence of $Ca^{2+}$. Data represents mean ± standard error from n = 3 technical replicates.

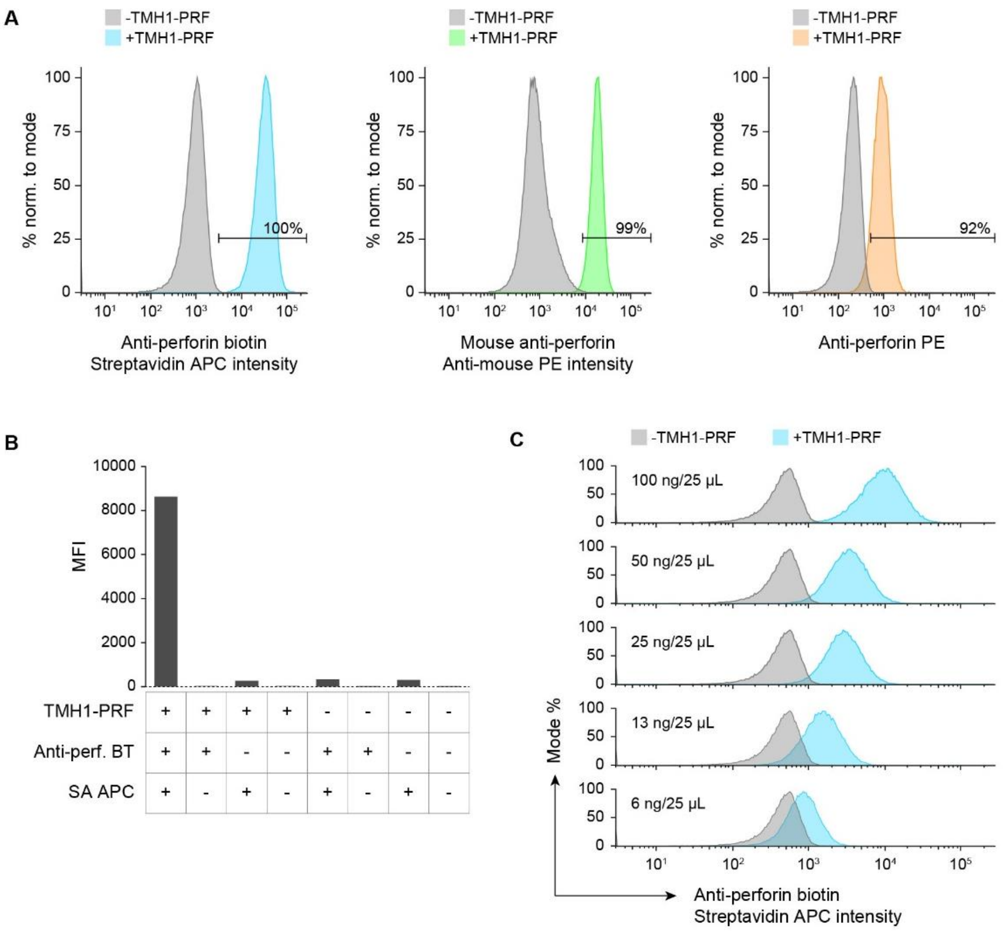


**Supplementary Figure 11: Anti-human perforin antibody reacts with disulphide-locked TMH1-PRF.**
**(A)** Three different conjugates of anti-human perforin antibodies (clone: δG9) from three different manufacturers (see **Supplementary Table 1**) produce fluorescent signals when TMH1-PRF is present on K562 cells, as detected by flow cytometry. All samples contained $10^5$ cells and were incubated with 100 ng TMH1-PRF for 15 min on ice and subsequently labelled in 1/25 antibody dilutions in 25 µL of media for 30 min on ice. Where applicable, primary and secondary stains were incubated sequentially.
**(B)** Using the biotinylated antibody shown in A), a substantial fluorescence signal was only obtained when TMH1-PRF, the biotinylated anti-perforin antibody (anti-perf. BT), and the streptavidin conjugated fluorophore (SA APC) were present.
**(C)** When antibody concentration and cell number were fixed, the brightness of the fluorescent signal depended on the concentration of TMH1-PRF, indicated in each panel.

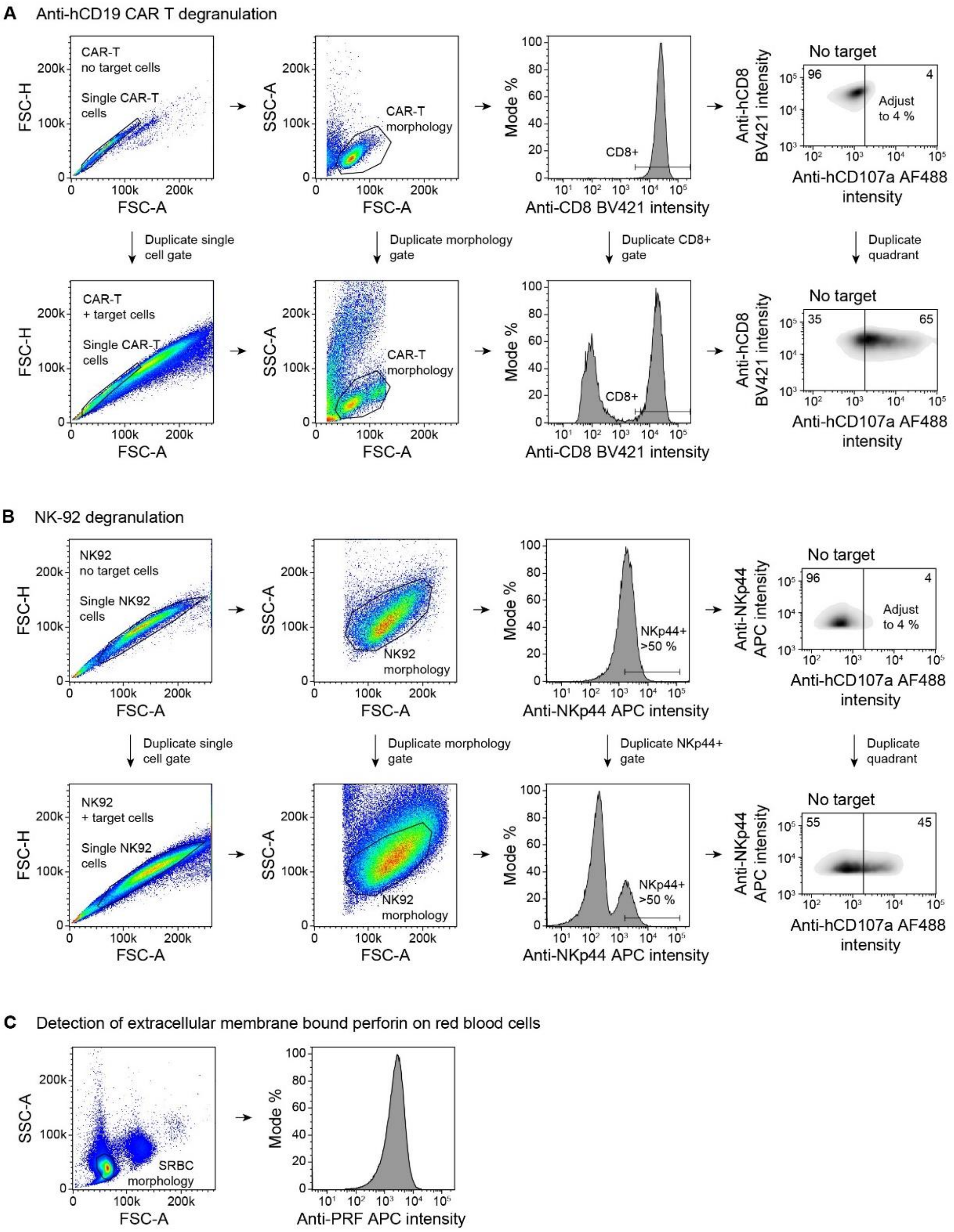


**Supplementary Figure 12: Analysis strategies for flow cytometry data.**
**(A)** Analysis strategy to evaluate degranulation of CAR T cells.
**(B)** as in A) for NK-92 cells.
**(C)** Morphology gating applied to SRBCs to evaluate perforin binding.

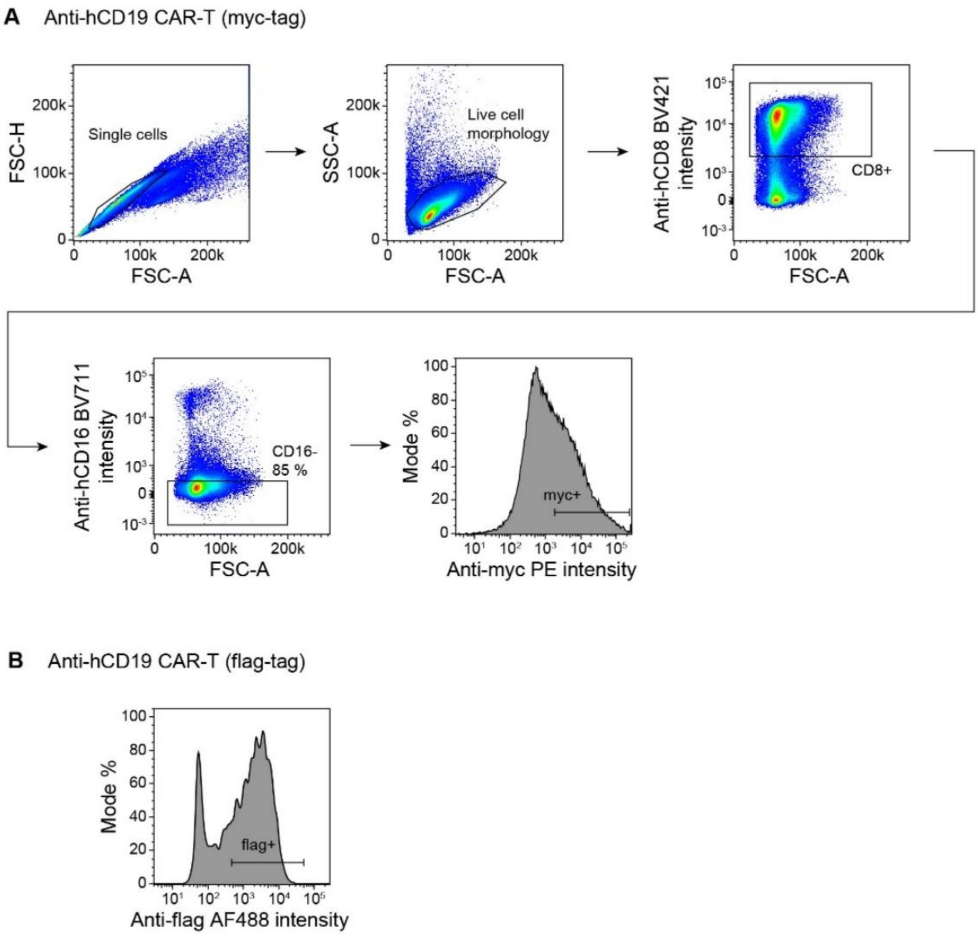


**Supplementary Figure 13: FACS strategy for CAR T cells.**
**(A)** Gating strategy to sort for retroviral vector transduced CAR T cells expressing an anti-hCD19 CAR with a myc-tag.
**(B)** Gating strategy to sort for lentiviral vector transduced CAR T cells was done as in A) except for using a flag-tag, with an example gating shown.

### Supplementary Tables

**Supplementary Table 1: Table of antibodies used in this study.**

| Name | Target | Reactivity | Host | Conjugated fluorophore | Clone | Manufacturer | Cat no. |
|---|---|---|---|---|---|---|---|
| Anti-hCD107a AF488 | CD107a | Human | Mouse | AF488 | H4A3 | BioLegend | 328610 |
| Anti-H-2Kb+SIINFEKL | SIINFEKL bound to H-2K$^b$ | Human | Mouse | PE | 25-D1.16 | BioLegend | 141603 |
| Anti-hCD56 PE | CD56 | Human | Mouse | PE | 5.1H11 | BioLegend | 362508 |
| Anti-hCD54 PE | CD54 | Human | Mouse | PE | HA58 | BD Biosciences | 555511 |
| Anti-hCD45r FITC | CD45RA | Human | Mouse | FITC | HI100 | BD Biosciences | 555488 |
| Anti-hNKp44 APC | NKp44 | Human | Mouse | APC | 253415 | R&D Systems | FAB22491A |
| Anti-hNKp44 PE | NKp44 | Human | Mouse | PE | 253415 | R&D Systems | FAB22491P |
| Anti-perforin PE | Perforin | Human/Mouse* | Mouse | PE | δG9 | BioLegend | 308106 |
| Anti-perforin | Perforin | Human/Mouse* | Mouse | - | δG9 | BD Biosciences | 556434 |
| Anti-perforin biotin | Perforin | Human/Mouse* | Mouse | - | δG9 | Ancell | 358-030 |
| Streptavidin APC | Biotin | - | Mouse | APC | - | eBioscience | 17-4317-82 |
| Anti-hCD8 BV421 | CD8 | Human | Mouse | BV421 | SK1 | BioLegend | 344748 |
| Anti-hCD16 BV711 | CD16 | Human | Mouse | BV711 | 3G8 | BioLegend | 302044 |
| Anti-hCD3 | CD3 | Human | Mouse | - | OKT3 | Invitrogen | 16-0037-85 |
| Anti-mouse IgG PE | Mouse IgG | Mouse | Goat | PE | Poly-clonal | Life Technologies | A10543 |
| Anti-flag AF488 | Flag-tag | - | Rat | AF488 | L5 | BioLegend | 637318 |
| Anti-myc | Myc-tag | - | Mouse | - | 9B11 | Cell Signaling Tech. | 2276 |
| Anti-mCD90.2 FITC | CD90.2 | Mouse | Rat | FITC | 53-2.1 | BD Biosciences | 553004 |
| Anti-mouse IgG PE | Mouse IgG | Mouse | Goat | PE | Poly-clonal | Life Technologies | A10543 |
| Anti-mCD28 | CD28 | Mouse | Hamster | - | 37.51 | BD Biosciences | 553295 |
| Anti-mCD3ε | CD3ε | Mouse | Hamster | - | 145-2C11 | BD Biosciences | 553058 |

* As reported in this manuscript

**Supplementary Table 2: Osmolarity measurements of pH stable media compared to standard cell media.**

| Medium | Osmolarity (mOsmol/kg)* |
|---|---|
| pH stable media pH 7.4 | 317±1 |
| pH stable media pH 6 | 304±2 |
| Human T cell medium | 289±3 |
| DMEM, 1X Glutamax, 10% FCS | 347±2 |
| 20 mM MES in Milli-Q $H_2O$ | 26±16 |
| 20 mM HEPES in Milli-Q $H_2O$ | 20±6 |

* Mean ± standard deviation from three technical replicates

**Supplementary Table 3: STR results and analysis for cell line identification.**

| Cell line | Amelogenin | CSF1PO | D13S317 | D16S539 | D21S11 | D5S818 | D7S820 | TH01 | TPOX | vWA | STR reference profile | Tanabe match |
|---|---|---|---|---|---|---|---|---|---|---|---|---|
| K562 | x, x | 9, 10 | 8, 8 | 11, 12 | 29, 30, 31 | 11, 12 | 9, 11 | 9.3, 9.3 | 8, 9 | 16, 16 | K-562 ATCC CCL-243 | 98% |
| U937 | X, X | 12, 12 | 10, 12 | 12, 12 | 27, 29 | 12, 12 | 9, 11 | 6, 9.3 | 8, 11 | 14, 15 | U937 ATCC CRL-1593.2 | 85% |
